# The characterization of multiple novel paramyxovirus species highlights the diverse nature of the subfamily *Orthoparamyxovirinae*

**DOI:** 10.1101/2022.03.10.483612

**Authors:** Bert Vanmechelen, Sien Meurs, Marie Horemans, Arne Loosen, Tibe Joly Maes, Lies Laenen, Valentijn Vergote, Fara Raymond Koundouno, N’faly Magassouba, Mandy Kader Konde, Ibrahima Sory Condé, Miles W. Carroll, Piet Maes

## Abstract

The subfamily *Orthoparamyxovirinae* is a group of single-stranded, negative-sense RNA viruses that contains many human, animal and zoonotic pathogens. While there are currently only 34 recognized member species in this subfamily, recent research has revealed that much of its diversity remains to be characterized. Using a newly developed nested PCR-based screening assay, we report here the discovery of fifteen orthoparamyxoviruses in rodents and shrews from Belgium and Guinea, thirteen of which are believed to represent new species. Using nanopore sequencing, complete genomes could be determined for almost all of these viruses, enabling a detailed evaluation of their genome characteristics. While most viruses are thought to belong to the rapidly expanding genus *Jeilongvirus*, we also identify novel members of the genera *Narmovirus, Henipavirus* and *Morbillivirus*. Together with other recently discovered orthoparamyxoviruses, both the henipaviruses and the morbillivirus discovered here appear to form distinct rodent-/shrew-borne clades within their respective genera, clustering separately from all currently classified member species. In the case of the henipaviruses, a comparison of the different members of this clade revealed the presence of a secondary conserved open reading frame, encoding for a transmembrane protein, within the F gene, the biological relevance of which remains to be established. While the characteristics of the viruses described here shed further light on the complex evolutionary origin of paramyxoviruses, they also illustrate that the diversity of this group of viruses in terms of genome organization appears to be much larger than previously assumed.

**Data availability:** The genome sequences generated in this study have been submitted to GenBank (accession numbers OK623353-OK623368).

## Introduction

In the past decade, the increased availability and reduced cost of next-generation sequencing (NGS) platforms have resulted in the discovery of countless novel virus species [1, 2]. A significant contribution to our understanding of viral diversity has been made by using large-scale metagenomics to expand the scope of putative host species. However, much of the virosphere also remains unexplored in previously established virus reservoirs, as demonstrated by the continuous discovery of new viruses in well-known virus hosts such as rodents and bats [3]. Although the pathological relevance and zoonotic potential of many of these newly discovered viruses is difficult to estimate solely from their genome sequence, valuable information can nonetheless be deduced by inferring their phylogenetic relationship with known viruses [4]. As illustrated by several recent large viral epidemics and pandemics (e.g. Ebola virus, Zika virus, SARS-CoV-2), the world is ill-prepared for adequately containing and managing virus outbreaks, especially if they are caused by unknown viruses (SARS-CoV-2) or viruses previously only reported in other parts of the world (Ebola virus, Zika virus) [5]. Through an improved understanding of the spread and diversity of animal-borne viruses, the risk of emergence of known or novel zoonoses locally or globally can be estimated more accurately, representing an important first step in improving global viral epidemic preparedness.

One virus family of particular interest for the potential emergence of novel human pathogens is the family *Paramyxoviridae*. Second only to the family *Rhabdoviridae*, paramyxoviruses form one of the largest known families of negative-sense, single-strand RNA viruses [6]. All paramyxoviruses share a similar genome organization, having a 15-21 kb genome that encodes six major genes, although members of the genus *Jeilongvirus* and some other species can have one or more additional genes of which the function remains poorly understood [7]. Currently, the family *Paramyxoviridae* contains 78 recognized member species, spread across four distinct subfamilies, the largest being the subfamily *Orthoparamyxovirinae* [6]. In addition to several important animal pathogens (e.g. Canine distemper virus, Fer-de-lance virus, Rinderpest virus), this subfamily is also home to human pathogenic viruses such as measles virus, mumps virus and parainfluenza viruses, as well as zoonotic pathogens, like Hendra virus and Nipah virus, that can infect both animals and humans [7, 8]. In recent years, many new orthoparamyxoviruses have been discovered and several of them are assumed to be pathogenic for animals or even humans. Especially the genus *Jeilongvirus* has seen a rapid expansion, with new member species being discovered primarily in rodents and bats, but also in carnivores and Eulipotyphla [9–18]. Furthermore, PCR-based screening studies have shown that much of the orthoparamyxovirus diversity remains to be properly characterized [19–31]. These studies are often performed using the screening assays developed by Tong and colleagues, which are capable of detecting paramyxoviruses belonging to a diverse range of genera [32].

In this study, we screened 95 small mammals from Belgium and Guinea, belonging to a variety of rodent and shrew species, for the presence of paramyxoviruses. For this screening, we developed a custom nested RT-PCR that was found to outperform the semi-nested assay of Tong et al. in terms of detectable diversity. Additionally, we attempted to obtain complete genome sequences for each detected putative species. In addition to Beilong virus (BeiV) and Pohorje myodes paramyxovirus 1 (PMPV-1), thirteen novel putative species were detected, belonging to the genera *Jeilongvirus, Morbillivirus, Narmovirus* and *Henipavirus*. For all but one of these novel viruses, a (near-) complete genome sequence could be determined.

## Materials and methods

### Sample collection

Cadavers of small mammals (rodents and shrews) that had been killed by cats or vehicles were collected in Belgium and stored at −20°C prior to transport to the laboratory. Following dissection, organs were stored in RNA*later* (Thermo Fisher Scientific, Waltham, MA, USA) at −80°C. In addition, organs were also collected from rodents and shrews that had been trapped and killed near Guéckédou (Nzérékoré Region, Guinea) in the frame of routine pest control. Identification of the caught animals was done by amplifying and sequencing part of the mitochondrial *cytochrome b* gene using the primer set (CytB Uni fw 5’-TCATCMTGATGAAAYTTYGG-3’, CytB Uni rev 5’-ACTGGYTGDCCBCCRATTCA-3’) developed by Schlegel and colleagues [33].

### RNA extraction

RNA was extracted from kidney tissue with or without viral enrichment procedures. Homogenization was performed by shredding 20 mg tissue in a tube filled with 135 μl sterile PBS (Thermo Fisher Scientific) and five zirconium oxide beads (2.8 mm, Thermo Fisher Scientific), using a Bertin homogenizer for two minutes at 4.000 rpm. For standard extractions, the homogenate was immediately subjected to RNA extraction using the RNeasy mini kit (Qiagen, Venlo, The Netherlands) according to the manufacturer’s instructions. For viral enrichment, 20X homemade enzyme buffer (1M Tris, 100 mM CaCl_2_ and 30 mM MgCl_2_, pH 8), 1 μl microccocal nuclease (New England Biolabs, Ipswich, MA, USA) and 2 μl benzonase (Merck-Millipore, Burlington, MA, USA) were added to the homogenate. After a two-hour incubation at 37°C, EDTA was added to a final concentration of 10 mM to stop enzymatic digestion. The homogenate was then centrifuged at 17.000g for three minutes, after which the resulting supernatant was filtered through a 0.8 μM filter by centrifuging at 17.000g for one minute. The resulting filtrate was then subjected to RNA extraction using the Viral RNA mini kit (Qiagen), according to the manufacturer’s instructions.

### Screening PCR

Nested RT-PCRs were performed using the OneStep RT-PCR kit (Qiagen). In each reaction, 2 μl RNA extract was mixed with 10 nmol dNTPs, 5X reaction buffer, 30 pmol forward primer (SCR_14F: 5’-ATGATGAARGGNCATGC-3’), 30 pmol reverse primer (SCR_26R: 5’-GCYTTRTCYTTCATRTACAT-3’) and 1 μl of the supplied enzyme mix, in a total volume of 25 μl. Cycling conditions were as follows: 30 minutes at 50°C and 15 minutes at 95°C, followed by 40 cycles of 30 seconds at 94°C, 30 seconds at 47°C and 1 minute at 72°C. A final elongation step was performed at 72°C for 10 minutes. For the inner reaction, 2 μl of the outer reaction product was used. All reaction and cycling conditions were identical to the ones used for the outer reaction, with the exception of the used primer set (SCR_16F: 5’-AARGGNCATGCHHTNTTCTG-3’, SCR_25R: 5’-TTCATRTACATDGTNAGRTC-3’) and the omission of the reverse transcription step at 50°C.

### Generation of cDNA libraries

RNA extracts from paramyxovirus-positive samples were converted into cDNA libraries using the SISPA method previously described by Greniger et al. [34]. Briefly, 4 μl RNA was reverse transcribed by adding 1 μl sol-PrimerA (40 pmol/μl, 5’-GTTTCCCACTGGAGGATA-N_9_-3’) and incubating at 65°C for five minutes and at 25°C for another five minutes. Next, 5 μl SuperScript IV master mix was added, consisting of 5x SSIV buffer, 10 nmol dNTPs, 50 nmol DTT, 20U RNaseOUT (Thermo Fisher Scientific) and 100U SuperScript IV Reverse Transcriptase (Thermo Fisher Scientific). The resulting mix was incubated at 23°C for 10 minutes, followed by 10 minutes at 50°C. For second strand synthesis, 5 μl Sequenase mix was added, containing 1 μl Sequenase buffer, 3.85 μl water and 0.15 μl Sequenase Version 2.0 DNA Polymerase (Thermo Fisher Scientific). Following an 8-minute incubation step at 37°C, a secondary Sequenase mix, containing 0.45 μl dilution buffer and 0.15 μl enzyme, was added, and the 8-minute incubation at 37°C was repeated. For amplification, 5 μl of the cDNA was mixed with LongAmp Taq 2X Master Mix (New England Biolabs) and 100 pmol Sol-PrimerB (5’-GTTTCCCACTGGAGGATA-3’) in a total volume of 50 μl. Cycling conditions were as follows: 2 minutes at 94°C, followed by 30 cycles of 30 seconds at 94°C, 45 seconds at 50°C and 1 minute at 65°C, and a final 10-minute elongation step at 65°C. Clean-up was performed using 0.4X AMPure XP beads (Beckman Coulter, Brea, CA, USA) according to the manufacturer’s instructions.

### Nanopore sequencing

Preparation of sequencing libraries was done using the SQK-LSK109 kit (Oxford Nanopore Technologies) according to the manufacturer’s protocol. When sequencing more than one sample per flow cell, the EXP-NBD104 and EXP-NDB114 kits were used for barcoding of individual samples. For each sample, 200 ng cDNA was used as input and ^~^120 fmol library was loaded onto the R.9.4.1 flow cell. Sequencing was performed on a MinION Mk1B or GridION. Basecalling (and demultiplexing) was performed using Guppy v3 and above. Porechop (github.com/rrwick/Porechop) was used to remove sol-Primer sequences added during cDNA generation. To identify reads corresponding to paramyxovirus sequences, a tblastx search of all reads against a custom BLAST database containing a representative sequence of each known orthoparamyxovirus species was done.

### Genome completion

If sufficient in number (>100), the subset of paramyxovirus reads was assembled into contigs using Canu v2.0 [35]. These contigs were polished using Medaka v0.9+ to achieve high quality sequences. To close remaining gaps in the sequence and to confirm the sequence of regions with low coverage, primer sets were designed based on the Medaka-polished contigs and used for the RT-PCR amplification of regions of interest using the OneStep RT-PCR kit (Qiagen) according to the manufacturer’s instructions. When insufficient reads were available, primers were designed based directly on the sequence of the individual nanopore reads. The resulting amplicons were purified using PureIT ExoZap PCR CleanUp (Ampliqon, Odense, Denmark) and sent to Macrogen Europe (Amsterdam, The Netherlands) for Sanger sequencing. The obtained chromatograms were corrected using Chromas v2.7 and, if applicable, joined with the Medaka contigs using Seqman v7.0.0. Genome ends were determined using the Roche 5’/3’race kit, 2nd generation (Hoffmann-La Roche, Basel, Switzerland) as described previously [13].

### Phylogenetic analysis

The NCBI Open Reading Frame Finder (ncbi.nlm.nih.gov/orffinder) was used to identify the open reading frames (ORFs) present in the newly sequenced genomes. For each of the six major proteins (N, P, M, F, G and L), separate alignments were made using mafft v7.489, incorporating a representative sequence of each (putative) paramyxovirus species for which a (near-) complete genome sequence is publicly available [36]. A list of all used accession numbers can be found in Supplementary Table S1. The resulting alignments were trimmed with trimAl v1.4.rev15, removing all columns with gaps in >10% of all sequences, after which the trimmed alignments were concatenated [37]. Following model selection using IQ-TREE, phylogenetic trees were inferred using Beast v1.10.4, employing an LG+G4+I amino acid substitution model [38–40]. The Beilong virus tree was made similarly but based on the complete nucleotide sequences of all available Beilong virus genomes, employing a GTR+G4+I nucleotide substitution model [41]. Analyses were run until adequate sample sizes were obtained (ESS>200) and maximum clade credibility trees were summarized from the posterior tree distribution using TreeAnnotator with a burn-in of 10%. FigTree v1.4.3 was used to visualize the resulting trees.

### Attempts at virus isolation

Tissue homogenate was made by shredding 30-50 mg kidney tissue in in a tube filled with 200 μl sterile PBS (Thermo Fisher Scientific) and five zirconium oxide beads (2.8 mm, Thermo Fisher Scientific), using a Bertin homogenizer for two minutes at 4.000 rpm. The homogenate was diluted 1/5 in Opti-MEM (Thermo Fisher Scientific) and centrifuged at 17.000g for two minutes to pellet cell debris. Vero E6 or BHK21J cells, grown to 80% confluency in 6-well plates, were then overlayed with 250 μl diluted homogenate for 90 minutes. Next, the homogenate was removed and cells were washed once with 1 ml sterile PBS before adding 2 ml DMEM supplemented with 2.5% Fetal bovine serum (Biowest), 200 mM L-glutamine, 1% Penicillin-Streptomycin, 1% Gentamicin and 0.2% Fungizone (all Thermo Fisher Scientific). Cells were inspected daily for cytopathogenic effects and passaged weekly three times. Supernatans from each passage was subjected to RNA extraction using the Viral RNA mini kit (Qiagen) and the resulting RNA was screened for the presence of paramyxovirus using the nested RT-PCR described above.

## Results

### Host determination and paramyxovirus screening

To investigate the presence of previously undiscovered paramyxoviruses in small mammals, we collected 95 rodents and shrews from sampling sites in Guinea (n=71) and Belgium (n=24) (Table 1). Using Sanger sequencing of the mitochondrial *Cytochrome b* gene, eighteen different host species could be identified. We initially screened the 71 Guinean animals for the presence of paramyxoviruses using an in-house developed nested RT-PCR. Twenty-two samples were found to be positive (31%). To validate whether there were any false negatives, we also screened these 71 animals using the ‘PAR’ and ‘RES-MOR-HEN’ primer sets previously developed by Tong et al. [32]. However, these assays showed a reduced performance compared to our newly developed primer set, detecting only six and nine positives, respectively. Both assays also failed to detect any additional positive samples. Therefore, the Belgian samples were only screened using our novel assay. An additional ten positives were found (41.7%), bringing the total to 32 positive samples, spread across twelve of the eighteen host species. Based on the sequences obtained by Sanger sequencing the screening amplicons, these 32 positives were thought to belong to fifteen separate paramyxovirus species. Putative species that were found in multiple animals were only detected in members of the same host species, and only in *Mastomys erythroleucus* and *Microtus agrestis-mice* did we detect more than one paramyxovirus species.

**Table 1:** Overview of collected animals

| Host species | Paramyxovirus<br>positivity rate<br>(Positives/total) | Country |
| --- | --- | --- |
| *Mastomys erythroleucus* | 11/42 | Guinea |
| *Mastomys natalensis* | 2/9 | Guinea |
| *Uranomys ruddi* | 4/6 | Guinea |
| *Crocidura grandiceps* | 1/3 | Guinea |
| *Lophuromys machangui* | 2/3 | Guinea |
| *Praomys rostratus* | 2/2 | Guinea |
| *Dasymys rufulus* | 0/2 | Guinea |
| *Hybomys planifrons* | 0/1 | Guinea |
| *Hylomyscus simus* | 0/1 | Guinea |
| *Mus setulosus* | 0/1 | Guinea |
| *Mus minutoides* | 0/1 | Guinea |
| *Crocidura russula* | 3/8 | Belgium |
| *Microtus arvalis* | 2/5 | Belgium |
| *Rattus norvegicus* | 1/4 | Belgium |
| *Apodemus sylvaticus* | 1/3 | Belgium |
| *Microtus agrestis* | 2/2 | Belgium |
| *Myodes glareolus* | 1/1 | Belgium |
| *Sorex minutus* | 0/1 | Belgium |

### Genome completion

To validate the hypothesis that the differences between the fifteen sequence groups were sufficient to warrant their classification as separate species, we attempted to determine the complete genome sequence for a representative strain from each group. Nanopore sequencing of randomly amplified cDNA was used to obtain more sequence information for each putative species, after which targeted RT-PCRs spanning remaining gaps or poorly covered regions of the genome were used to determine the rest of the genome sequence. Genomes were completed by Sanger sequencing of the genome ends following 5’/3’ RACE. Through this approach, (near-) complete genomes could be obtained for thirteen of the fifteen putative species. For a fourteenth sequence, the 3’ end of the genome and one gap covering part of the G and L ORFs could not be determined due to the low integrity of the available RNA. Lastly, repeated efforts to obtain additional sequence information for the fifteenth virus, which was found in three Guinean *Mastomys erythroleucus*-samples, only managed to identify two small fragments of the N and L ORFs, presumable due to a very low viral load in these three samples. An overview of the organization of the different obtained genomes and some of their characteristics are shown in Figure 1 and Table 2. The naming of the different viruses is based on the genus to which they belong, the host in which they were detected, the location in which these hosts were found or a combination of these characteristics.

**Figure 1:**
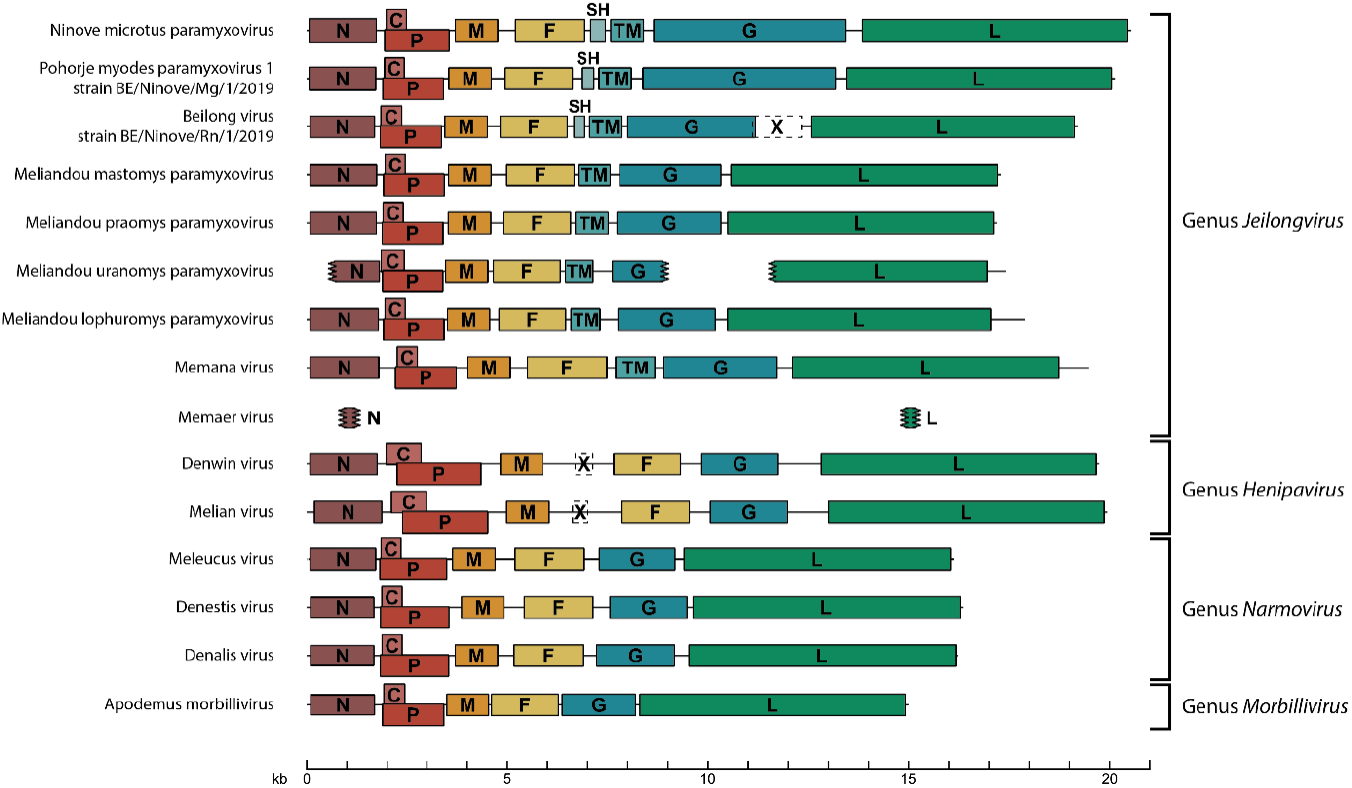
Genome organization of novel paramyxoviruses. The fifteen viruses described in this study have differing genome layouts, which seem to correlate with their presumptive classification in four separate genera. All genomes and ORFs they encode are drawn according to scale. ‘X’ ORFs indicate ORFs of significant size (>100 amino acids) of which it is unclear if they are biologically active.

**Table 2:** Overview of detected paramyxoviruses

| Genbank accession | Virus | Host species | Country | Complete (Y/N) | Putative genus |
| --- | --- | --- | --- | --- | --- |
| OK623353 | Melian virus | *Crocidura grandiceps* | Guinea | Y | *Henipavirus* |
| OK623354 | Denwin virus | *Crocidura russula* | Belgium | Y | *Henipavirus* |
| OK623355 | Ninove microtus paramyxovirus | *Microtus agrestis* | Belgium | Y | *Jeilongvirus* |
| OK623356 | Apodemus morbillivirus | *Apodemus sylvaticus* | Belgium | Y | *Morbillivirus* |
| OK623357 | Pohorje myodes paramyxovirus 1 – strain<br>BE/Ninove/Mg/1/2019 | *Myodes glareolus* | Belgium | Y | *Jeilongvirus* |
| OK623358 | Beilong virus – strain BE/Ninove/Rn/1/2019 | *Rattus norvegicus* | Belgium | Y | *Jeilongvirus* |
| OK623359 | Denestis virus | *Microtus agrestis* | Belgium | Y | *Narmovirus* |
| OK623360 | Denalis virus | *Microtus arvalis* | Belgium | Y | *Narmovirus* |
| OK623361 | Meliandou mastomys paramyxovirus | *Mastomys erythroleucus* | Guinea | Y | *Jeilongvirus* |
| OK623362 | Meliandou praomys paramyxovirus | *Praomys rostratus* | Guinea | Y | *Jeilongvirus* |
| OK623363 | Memana virus | *Mastomys natalensis* | Guinea | Y | *Jeilongvirus* |
| OK623364 | Meliandou lophuromys paramyxovirus | *Lophuromys machangui* | Guinea | Y | *Jeilongvirus* |
| OK623365 | Meleucus virus | *Mastomys erythroleucus* | Guinea | N | *Narmovirus* |
| OK623366 | Meliandou uranomys paramyxovirus | *Uranomys ruddi* | Guinea | N | *Jeilongvirus* |
| OK623367 - OK623368 | Memaer virus | *Mastomys erythroleucus* | Guinea | N | *Jeilongvirus* |

Excluding the three sequences for which part of the genome sequence remains to be determined, the genomes sequenced here are all compliant with the ‘rule-of-six’, which states that paramyxovirus genomes must have a length that is an exact multiple of six to enable genome replication [42]. However, their exact lengths vary greatly, representing both the smallest (14,994 nucleotides) and largest (20,454 nucleotides) known orthoparamyxoviruses. The diverse nature of the genomes characterized in this study is further illustrated by their scattered phylogenetic clustering throughout the entire subfamily *Orthoparamyxovirinae*, with the exception of a single clade that contains the genus *Respirovirus* and the non-mammalian genera *Ferlavirus* and *Aquaparamyxovirus* (Figure 2). While most viruses we found seem to belong to the genus *Jeilongvirus*, we also detected three novel narmoviruses, two henipaviruses and one morbillivirus. Unfortunately, attempts to rescue infectious virus from animal tissue using Vero E6 and BHK21J cells were unsuccessful.

**Figure 2:**
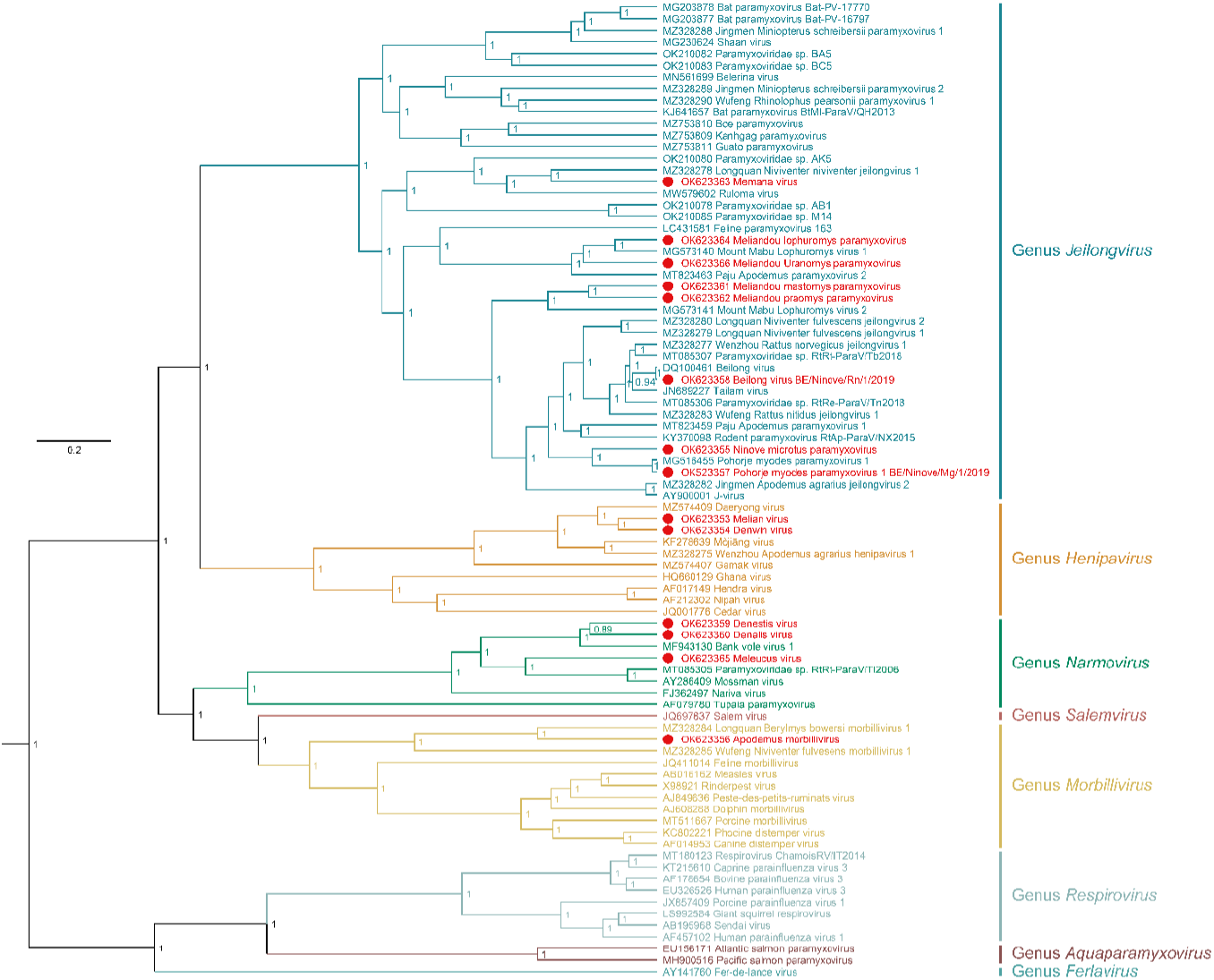
The subfamily *Orthoparamyxovirinae*. Bayesian phylogenetic tree of all 84 currently known members of the subfamily *Orthoparamyxovirinae* for which near-complete genomes are available. The tree was made based on a concatenation of individual alignments of the N, P, M, F, G and L ORFs, covering 4,422 sites after trimming. Missing sites in partial alignments were marked as such (‘?’) prior to phylogenetic inference. Different genera within the subfamily are marked in different colors, according to the current ICTV classification. New sequences presented here are marked in red and are preceded by a red dot. Values at the nodes indicate posterior support. Phylogenetic tree drawn according to scale, with the scale bar denoting the number of substitutions/site. Used accession numbers are shown in the tree. Memaer virus was not included due to a lack of available sequence information.

### Characterization of novel jeilongviruses

Nine of the fifteen paramyxoviruses detected here are thought to belong to the genus *Jeilongvirus*. This includes Memaer virus, for which insufficient sequence data could be obtained to do a meaningful phylogenetic analysis. Based on blastn comparisons, this virus appears to be most similar to the here reported Memana virus, Ruloma virus and Longquan Niviventer niviventer jeilongvirus 1 (Table 3), although sufficiently dissimilar, especially given the conserved nature of the detected regions, to consider Memaer virus a separate viral species. Conversely, two of the jeilongviruses detected in this study, Pohorje myodes paramyxovirus 1 (PMPV-1) strain BE/Ninove/Mg/1/2019 and Beilong virus (BeiV) strain BE/Ninove/Rn/1/2019, appear to fall within the already recognized species *Myodes jeilongvirus* and *Beilong jeilongvirus*, respectively. While the latter is intriguingly similar to all known BeiV genomes, phylogenetically clustering amidst them and sharing 94.50-96.14% nucleotide identity (Figure 3), the PMPV-1 sequence is more dissimilar from the only other known sequence of PMPV-1, sharing only 84.25% identity on nucleotide level. Interestingly, the variation between these two sequences appears to be roughly equal throughout the coding regions of the genome, with a notable exception being the C-terminal domain of the G protein (Figure 4). While the first half and the last quarter of this protein display a comparable degree of conservation as the rest of the genome, the third quarter bears little resemblance to the other available PMPV-1 sequence and appears to be even less conserved than the non-coding regions of the genome. A similar pattern, albeit less pronounced, can also be observed when comparing the BeiV sequence presented here with all known BeiV genomes.

**Table 3:**
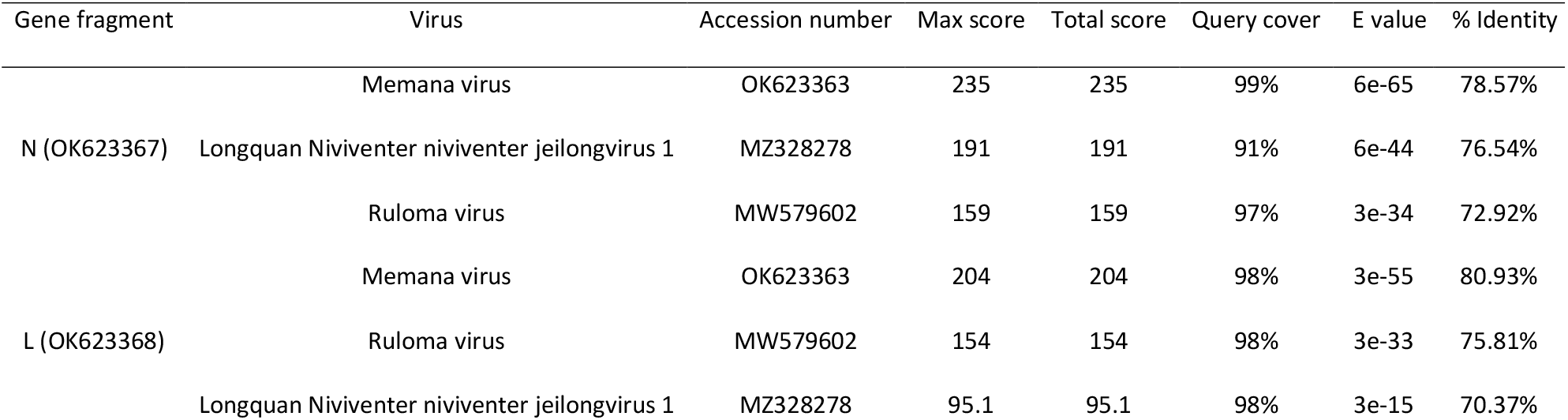
Blastn comparison of Memaer virus

**Figure 3:**
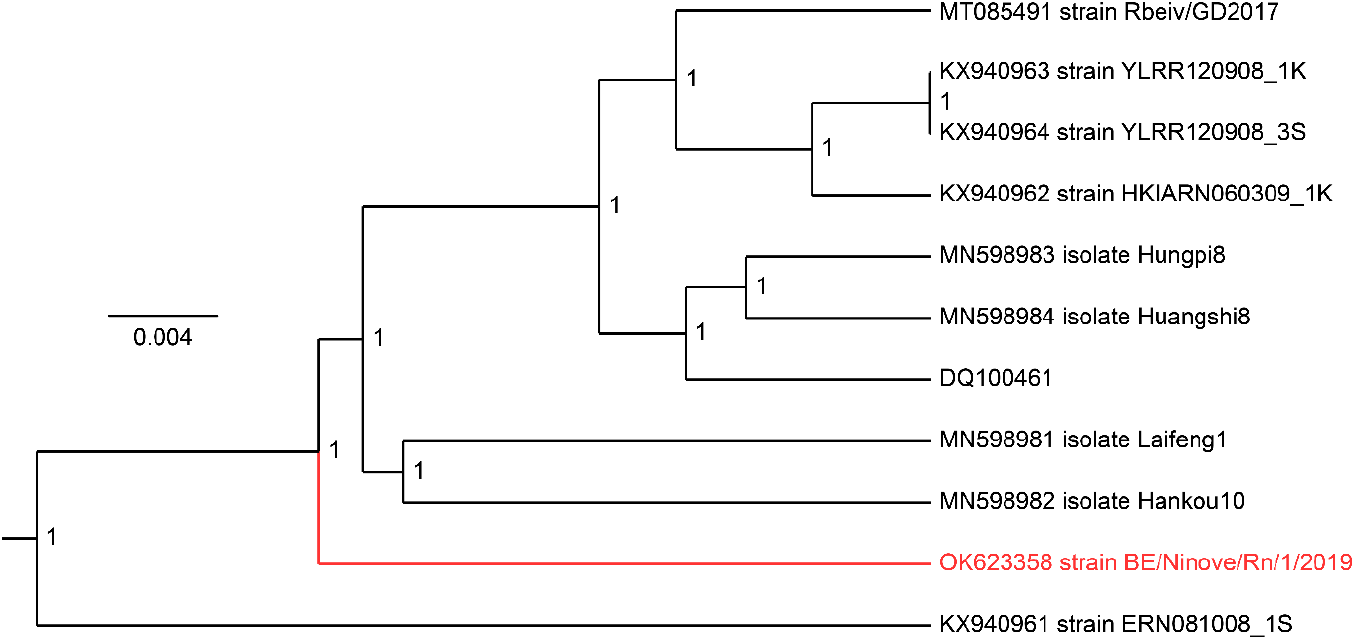
Phylogenetic tree of all near-complete Beilong virus genomes. Bayesian phylogenetic tree based on a nucleotide alignment of all available Beilong virus genome sequences. Despite its different geographical origin, the strain identified here (marked in red) clusters amongst the other known Beilong virus sequences from China and Hong Kong. Values at the nodes indicate posterior support. Phylogenetic tree drawn according to scale, with the scale bar denoting the number of substitutions/site. Used accession numbers are shown in the tree.

**Figure 4:**
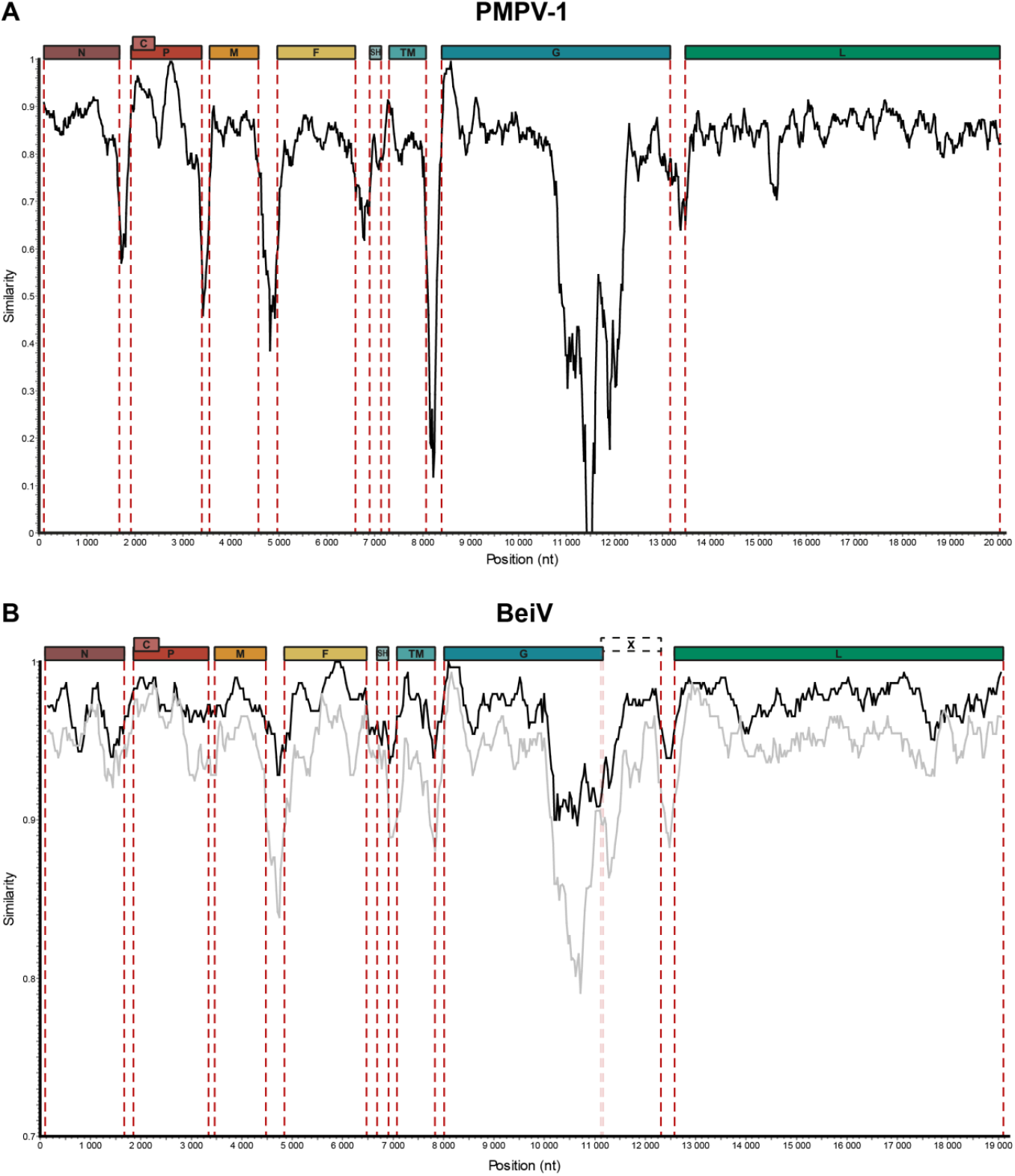
Similarity comparison of the new PMPV-1 and BeiV strains. Panel A shows a comparison of the two known PMPV-1 strains, highlighting a comparable degree of conservation all throughout the coding regions of the genome, with a notable exception in the middle of the G gene. Gene borders are marked by dashed red lines. Figure made using SimPlot v3.5.1, employing a window size of 200 bp and a step size of 20 bp. Panel B shows a comparable but significantly weaker pattern when comparing the new BeiV strain described here with all (black line) known BeiV genomes or only the most divergent one (KX940961; grey line). Gene borders are marked by dashed red lines, with faint lines indicating the transition from the G to the X ORF within the G gene. Figure made using SimPlot v3.5.1, employing a window size of 300 bp and a step size of 30 bp.

Previously, we showed that the glycoproteins of jeilongviruses share certain unique features not found in those of other paramyxoviruses [16]. In addition to their exceptional and highly variable length, the C-terminal regions of jeilongvirus G proteins are also characterized by a remarkably high fraction of proline, serine and threonine, despite their lack of sequence conservation at the nucleotide level. However, since our initial analysis, many new jeilongviruses have been discovered. Redoing the analysis discussed in [16], using all currently available jeilongvirus G genes, shows that the genus *Jeilongvirus* can be divided into five clades based on the length and sequence composition of the G protein (Figure 5). Members of clade I show no elongation and have G proteins that are comparable to those of other orthoparamyxoviruses. Members of clades II, III and IV show a modest increase in size and their extension is often but not necessarily characterized by P/T/S-enrichment. Lastly, members of clade V have exceptionally large G proteins, 2-3 times the average size of an orthoparamyxovirus G protein, and their proteins are all characterized by P/T/S-enrichment. Interestingly, several members of this clade have acquired stop codons in their G genes, resulting in the split of the original G ORF into two smaller ORFs, G and X. It remains to be determined if these X ORFs, which fall within the G gene are expressed, either independently or as part of the G protein. Additionally, while many seem to follow this same pattern, P/T/S-enrichment appears less ubiquitous than previously assumed (Figure 5). Interestingly, the peak in P/T/S-enrichment appears to coincide with the least conserved region at the nucleotide level, as illustrated by comparing Figures 4 and 5 for the novel strains of PMPV-1 and BeiV reported here.

**Figure 5:**
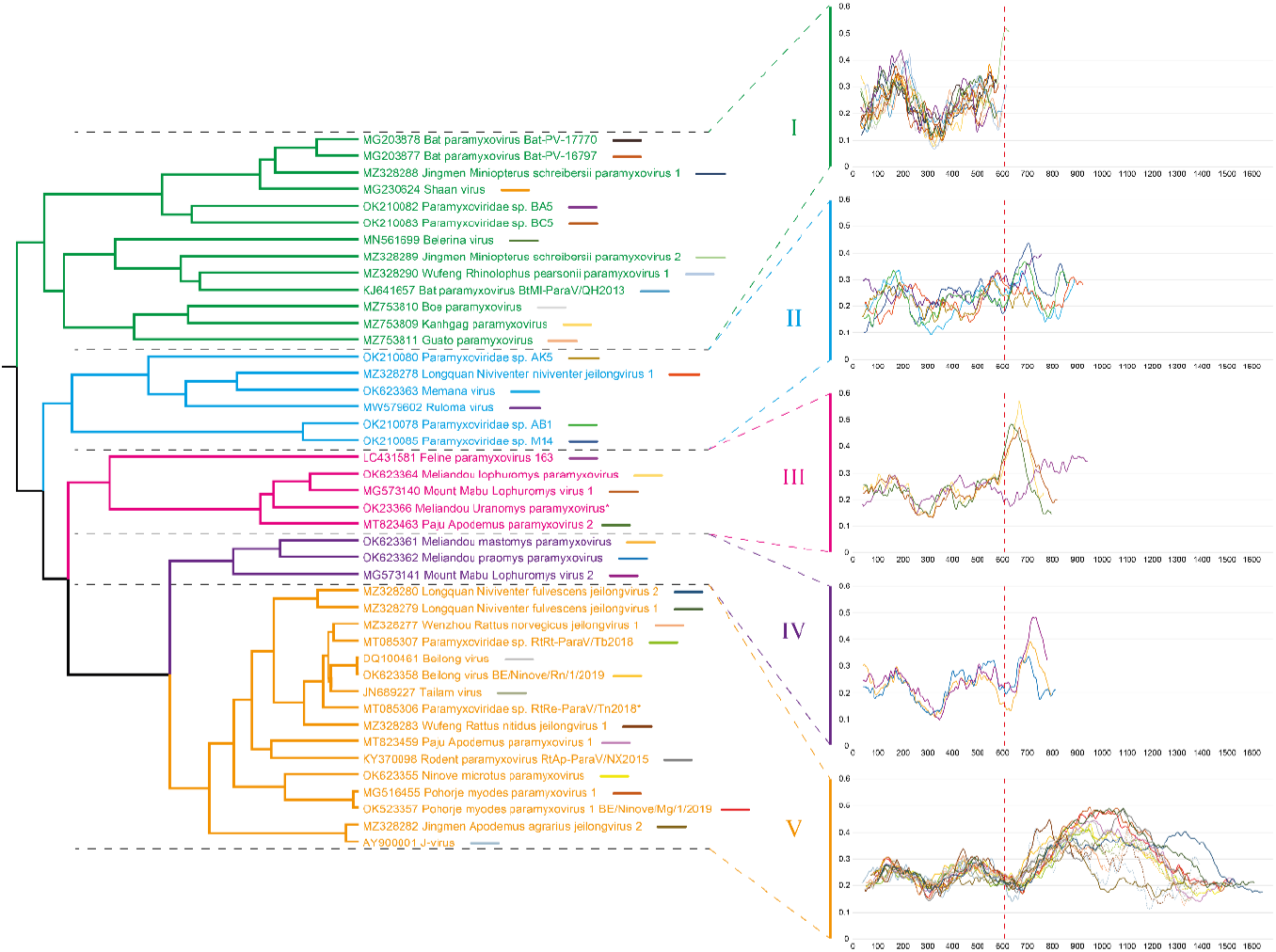
Different G protein organizations in the genus *Jeilongvirus*. Close-up of the tree shown in Figure 2, depicting only the genus *Jeilongvirus*. Next to the tree are graphs indicating the length of the different jeilongvirus G proteins and their cumulative fraction of proline, serine and threonine, calculated using a sliding window corresponding to 10% of the length of each protein. Based on their phylogenetic relationship and the organization of their G protein, jeilongviruses can be divided into five clades. Members of clade I have G proteins that are comparable in size to those of non-jeilongvirus orthoparamyxoviruses (average size: 607 nt, dashed red line). Conversely, the G proteins of members of clade V are up to 150% larger and their C-terminal extensions are characterized by a very high fraction of proline, serine and threonine. In some members of this clade, the G ORF has acquired a stop codon somewhere within this region, resulting in a secondary ‘X’ ORF within the G gene. It is unclear if this ‘X’ ORF is expressed, either by itself or as part of the G protein (dashed lines). Clades II, III and IV are characterized by a more modest expansion of the G protein (up to 50%) that is in some cases characterized by proline-, serine- and threonine-enrichment. Sequences marked by a ‘*’ are not included in the graphs because their G genes have not been fully sequenced.

### A new hypothetical protein characterizes a separate henipavirus clade

While most novel viruses described here were found in rodents, we also detected two new viruses in shrews: Denwin virus in a *Crocidura russula* or greater white-toothed shrew and Melian virus in a *Crocidure grandiceps* or large-headed forest shrew. Both of these viruses cluster within the genus *Henipavirus*, together with the recently discovered Gamak virus and Daeryong virus, who were also detected in shrews, and the rodent-borne Mojiang virus and Wenzhou Apodemus agrarius henipavirus 1 (Figure 2). Together, these viruses seem to form a distinct lineage within the genus *Henipavirus*, separate from all bat-borne henipaviruses. Interestingly, all the viruses within this clade, including Denwin virus and Melian virus, contain a small ORF (‘X’) between the M and F ORFs that is not seen in the genomes of other paramyxoviruses. This ORF seems to be conserved both at the nucleotide and amino acid level, coding for a small transmembrane protein (Figure 6). However, this ORF does not have its own gene start and stop signals but instead falls within the boundaries of the F gene. Whether this ORF is expressed as part of a multicistronic mRNA or through another mechanism remains to be established.

**Figure 6.**
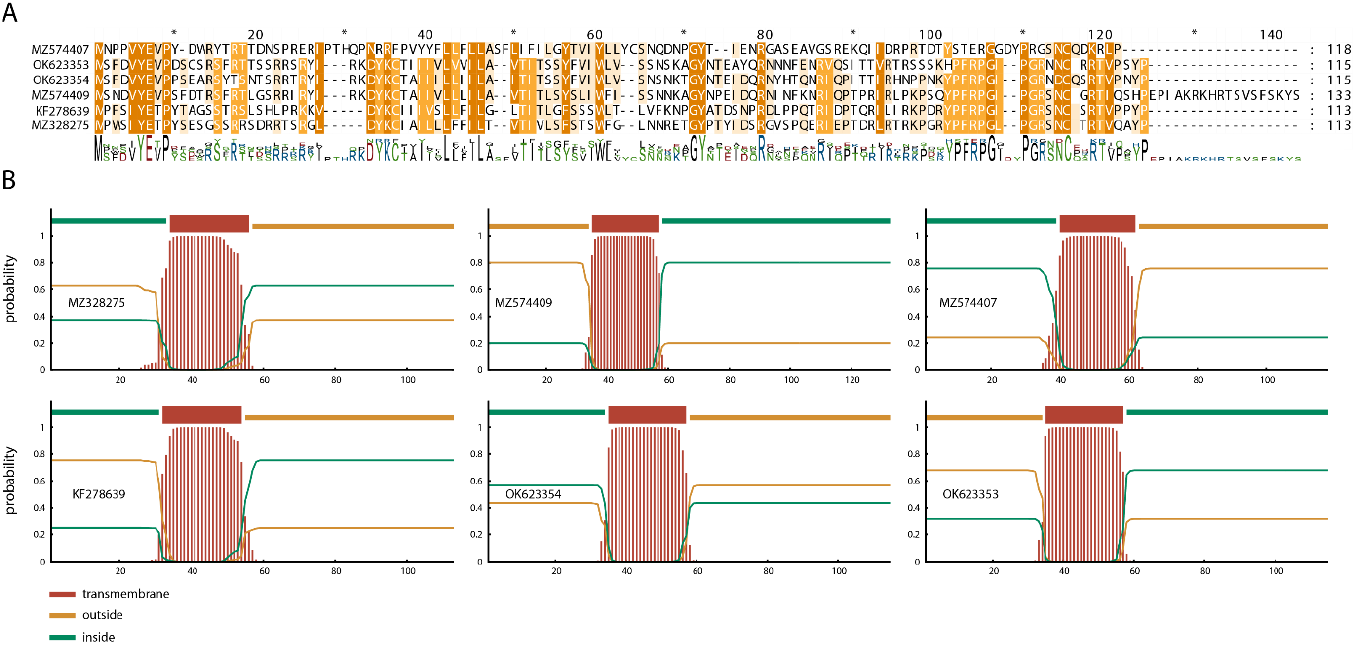
Conservation of a secondary ORF within the henipavirus F gene. **(A)** All members of the rodent-/shrew-borne clade of henipaviruses have a secondary ORF in the F gene, preceding the actual F ORF. Both the length and sequence composition of this ‘X’ ORF show a certain degree of conservation, hinting at the biological relevance of this ORF. **(B)** All these ‘X’ ORFs are predicted to encode a transmembrane protein. Plots made using TMHMM – 2.0 (https://services.healthtech.dtu.dk/service.php?TMHMM-2.0).

### Discovery of other paramyxoviruses

Besides jeilongviruses and henipaviruses, we also discovered three new narmoviruses and one morbillivirus. Unlike the abovementioned viruses, these four viruses have a more traditional paramyxovirus genome organization, characterized by only six genes (N-P/V/C-M-F-G-L) (Figure 1). In accordance with the hosts in which they were found, Denestis virus (Microtus agrestis) and Denalis virus (Microtus arvalis) cluster closely together, near Bank vole virus 1, while Meleucus virus is more closely related to Mossman virus and the yet unnamed ‘Paramyxoviridae sp. isolate RtRt-ParaV/Tl2006’ (Figure 2). Similar to the abovementioned situation in the genus *Henipavirus*, the clustering of Apodemus morbillivirus, alongside the recently discovered Wufeng Niviventer fulvescens morbillivirus 1 and Longquan Berylmys bowersi morbillivirus 1, appears to indicate the existence of a distinct rodent-borne morbillivirus clade, forming a distant sister clade to all non-rodent morbilliviruses.

## Discussion

Many novel orthoparamyxovirus species have been discovered in recent years, especially thanks to the increased availability of NGS platforms. However, despite the advancements that have been made, the cost per sample remains too high to allow the unfiltered sequencing of large sample collections [4]. In studies focusing on one or more particular pathogens, targeted methods that allow the selection of a specific subset of samples therefore remain of interest to maximize the return on investment of NGS experiments. Conversely, too stringent selection methods might result in interesting samples erroneously being excluded from further analysis. Therefore, we sought to make a PCR-based screening approach that would maximize the number and diversity of orthoparamyxoviruses that could be detected using a singular assay. By opting for a nested PCR approach that targets a small region of the L gene using four degenerate primers, we were able to detect a wide range of orthoparamyxoviruses, covering each major clade within the subfamily besides the one containing the genus *Respirovirus* and related non-mammalian viruses. Using our new assay, we found >33% of the animals in this study to be positive for paramyxoviruses. Even though this study had a limited sample size and the possibility of false negatives due to insufficient viral load or a too divergent sequence cannot be excluded, this percentage seems to fall in line with what has previously been reported in paramyxovirus screening studies of small mammals [18, 30, 43]. However, few previous studies have reported a similarly broad diversity as observed here, which might be attributable to the newly designed primer sets used in this study, as most previous studies were performed using the semi-nested assays developed by Tong and colleagues. Despite their supposed broader detection range, these assays failed to correctly identify most of the positive samples in our collection, potentially indicating that previous studies might have underestimated the true diversity of paramyxoviruses and that these viruses are even more diverse and more ubiquitously present than previously known.

The majority of the viruses we detected belong to the genus *Jeilongvirus*. Prior to the establishment of this genus in 2019, only six putative jeilongviruses genome sequences were known, although some studies had already hinted at the existence of many more viruses remaining to be characterized [31, 44, 45]. However, since 2019, this genus has known the strongest expansion of all paramyxovirus genera. Although the International Committee for the Taxonomy of Viruses (ICTV) currently only recognizes seven species in this genus, more than thirty additional putative species have been described and/or published on GenBank in the last three years and jeilongviruses now represent half of all known orthoparamyxoviruses (Figure 2, Figure 5) [6]. Interestingly, this expansion has also revealed a remarkable diversity in genome organization within members of this genus. Aside from some minor exceptions, paramyxovirus genomes contain six genes (N-P/V/C-M-F-G-L) and members of the same genus typically have comparable genome lengths [7]. Jeilongvirus genomes, however, contain one or two additional genes and can vary in length from <16 kb to >20.5 kb. The first of these extra genes is commonly named ‘TM’ because it encodes a protein with a transmembrane region. A protein like this is present in the genome of all jeilongviruses and, in the case of J-virus, it has been shown to interact with the F and G proteins and to promote cell-to-cell fusion [46]. However, even though all Jeilongviruses encode a TM protein, it remains to be established whether these proteins exert similar functions, as the sequences of TM proteins from different clades appear to be phylogenetically unrelated, differing strongly in terms of length and sequence composition. As noted previously (see [16]), a similar observation can be made for the second additional jeilongvirus gene, SH, which is present only in members of certain clades of the genus. Future research will be needed to determine whether the TM and SH proteins of different clades are functional homologues of each other and to determine the precise function of these proteins.

Like the genus *Jeilongvirus*, the genus *Narmovirus* was only established in 2019 and little is known about the pathological relevance of these viruses for animal and human health. Conversely, the genera *Morbillivirus* and *Henipavirus* were established many years ago and many of their members are known to be important human, animal or zoonotic pathogens [47, 48]. The new viruses described here, in combination with other novel genomes published by other groups in the last year, illustrate the existence of a separate clade of rodent- and/or shrew-borne viruses that diverged early on from the rest of the genus within both the genus *Henipavirus* and the genus *Morbillivirus*. A question that remains to be resolved is whether the members of these separate clades are equally pathogenic as their sister clade counterparts and, especially in the case of the henipaviruses, if they have the same zoonotic potential. A possible argument against this hypothesis might be that Mòjiāng virus, the first rodent-borne henipavirus to be discovered, has been shown to be incapable of using the same host-entry pathways as other henipaviruses due to antigenic differences in its G protein [49, 50]. Interestingly, the availability of additional sequences of related viruses has made it apparent that members of this rodent-/shrew-borne henipavirus clade contain a conserved ORF coding for a transmembrane protein nested within the F gene. Although the mechanism through which this ORF is expressed remains to be established, as well as the function of the resulting protein, its existence highlights that more complex genome organizations are not a unique feature of jeilongviruses but seem to be the result of independent host-specific adaptation events that have occurred in multiple lineages within the subfamily *Orthoparamyxovirinae*. It will be of interest to determine whether this transmembrane protein is a functional homologue to any of the transmembrane proteins found in jeilongviruses and if similar proteins can also be found in other paramyxovirus lineages that remain to be discovered.

## Acknowledgments

The authors wish to thank the prefectural health authorities of Guéckédou (DPS) for on-site assistance. B.V. was supported by a FWO SB grant for strategic basic research of the “Fonds Wetenschappelijk Onderzoek”/Research foundation Flanders (1S28617N).

## Author contributions

B.V. and P.M. designed the study. B.V., S.M., M.H., A.L. and T.J.M. performed the experimental work. L.L., V.V., M.W.C., F.R.K., I.S.C., M.K.K. and P.M. ensured sample collection. M.W.C. and P.M. provided reagents and materials. B.V. wrote the main manuscript text and prepared the figures. All authors reviewed the manuscript and approved the final version.

## Supplementary data

Supplementary Table S1: Overview of all used accession numbers

